# Faster relaxation of nonphotochemical quenching (NPQ) in C4 than in C3 species

**DOI:** 10.1101/2025.04.01.646649

**Authors:** Lucía Arce Cubas, Asuka Nakamura, Richard L. Vath, Julia Walter, Cristina Rodrigues Gabriel Sales, Emmanuel L. Bernardo, Yuri Nakajima Munekage, Johannes Kromdijk

## Abstract

Acceleration of photoprotective non-photochemical quenching (NPQ) responses to changes in light intensity has been suggested as a strategy to enhance crop yield. Despite many key crops utilising C4 photosynthesis, our current understanding of NPQ overwhelmingly comes from C3 species. Using a series of experiments on three phylogenetically controlled C3 and C4 comparisons, we show that NPQ relaxation is faster in C4 species. Temporal analysis of NPQ relaxation in leaves infiltrated with inhibitors to block proton motive force formation or xanthophyll de-epoxidation showed that the faster relaxation observed in C4 species is driven by a greater contribution of energy-dependent quenching (qE) to overall NPQ. We show that the C4-associated enhancement of qE is linked to altered regulation of lumen pH in C4 species, reflecting increases in cyclic electron flow and membrane proton conductivity to meet the increased ATP demands of the C4 pathway. Indeed, in two of the three tested C4 species, NPQ relaxation became significantly slower and statistically indistinguishable from paired C3 species when ATP and NADPH consumption was suppressed by performing measurements in CO_2_-free air. Altogether, our results suggest that NPQ responses in C4 species may already be optimised to maintain high photosynthetic efficiency in the fluctuating light conditions typically found within C4 canopies. Given the intrinsically faster NPQ in C4 photosynthesis, further acceleration of NPQ may have limited scope to enhance crop photosynthetic efficiency.

**Significance Statement:** Acceleration of non-photochemical quenching has been proposed as a means to enhance crop photosynthetic efficiency in C3 species but whether this strategy has potential in C4 species, which include several major crops, remains unclear. We use three phylogenetically paired C3 and C4 species to show that NPQ relaxation is significantly faster in species with the C4 pathway, possibly aiding the maintenance of photosynthetic efficiency in fluctuating light environments. As a result, accelerating the rate of NPQ relaxation in C4 crops may have a more limited scope to enhance photosynthesis.

## Introduction

Nonphotochemical quenching (NPQ) refers to a collection of photoprotective mechanisms wherein excess light energy in photosystem II (PSII) is dissipated as heat, preventing overexcitation and the formation of reactive oxygen species that would otherwise damage the photosynthetic machinery (1). NPQ components operate at different timescales and likely involve conformational changes in PSII-associated antennae that trigger the quenched state (2). Energy-dependent quenching (qE) is activated within seconds to minutes by lumen acidification (3), which also triggers xanthophyll cycle enzyme violaxanthin de-epoxidase (VDE) to convert violaxanthin to zeaxanthin, enhancing qE (4). Zeaxanthin accumulation further supports a qE-independent quenching component (qZ) that activates and recovers over minutes to hours (5). Even more sustained quenching (qH) relies on the plastid lipocalin LCNP and is negatively regulated by suppressor of quenching SOQ1 (6). Finally, photoinhibitory quenching (qI) is associated with photodamage and requires *de novo* synthesis of the D1 protein for recovery (7).

Our molecular understanding of NPQ as detailed above overwhelmingly comes from C3 species, where NPQ relaxation has been found to significantly lag behind changes in irradiance, temporarily lowering photosynthetic efficiency and leading to substantial losses in ‘foregone’ canopy carbon assimilation (8). Accelerating recovery from photoprotection represents a promising strategy for improving crop yield in C3 species (9, 10), yet whilst many key crops are C4 (11) the specifics of the C4 NPQ response remain largely unknown (12).

In C3 photosynthesis, CO_2_ is directly fixed in mesophyll (M) chloroplasts by ribulose-1,5-biphosphate carboxylase/oxygenase (Rubisco) into 3-carbon compound 3-phosphoglycerate (3-PGA). Higher photosynthetic rates are generally found in C4 species, where a carbon concentrating mechanism (CCM) enhances photosynthesis by suppressing RuBP oxygenation and concomitant photorespiration (13). The C4 pathway operates between morphologically distinct M and bundle sheath (BS) cells, typically arranged in ‘Kranz’ anatomy: CO_2_ is initially converted to bicarbonate in the M and fixed into 4-carbon oxaloacetate that is further reduced or transaminated into malate or aspartate before diffusion into the BS, where decarboxylating enzymes release CO_2_ around Rubisco and into the C3 cycle (14). Different C4 “subtypes” use different decarboxylases, often in combination (15), and have additional cell and subtype-specific ATP requirements for the regeneration of CCM biochemical intermediates (16-18). The ATP:NADPH ratio generated by linear electron flow (LEF) from PSII to PSI is insufficient to satisfy the demands of the C3 cycle, and the additional ATP demands by C4 metabolism further the imbalance (19). Cyclic electron flow (CEF) helps balance energy budgets by recycling electrons around PSI back to plastoquinone (PQ), contributing to the proton motive force (*pmf*) that powers ATP synthesis without concurrent production of NADPH (20). Considerably higher ratios of PSI:PSII (21, 22) and CEF:LEF (23) are found in C4 versus C3 species, reflecting the ATP cost of the C4 pathway.

The functional differences between C3 and C4 photosynthesis are likely to affect NPQ. In C3 species, CEF plays a major photoprotective role by contributing to ΔpH and thus qE activation, with CEF-defective mutants having severely reduced NPQ (24, 25). This could suggest an enhanced qE component in C4 species, given their higher CEF:LEF ratios. In C3 species, CEF is predominantly mediated by proton gradient regulation 5 (PGR5) and PGR5-like photosynthetic phenotype 1 (PGRL1), with a minor contribution by chloroplast NADH dehydrogenase-like complex (NDH) in low and fluctuating light (26-28). In C4 species, although PGR5/PGRL1 is still important, recent studies show CEF occurs primarily via NDH (29-31). Notably, NPQ amplitude was *lower* in C4 PGR5 and PGRL1-deficient mutants, but *higher* in NDH mutants (30, 31). Based on these results, it was suggested that PGR5/PGRL1 could play a similar photoprotective role in C4 photosynthesis as in C3 species, with NDH mainly contributing to supplementing ATP production (31), but the effect of CEF pathway interplay on C4 NPQ remains unclear. Finally, NPQ modulation is also regulated by the proton conductivity of the thylakoid membrane (gH^+^, indicative of ATP synthase activity), as changes to gH^+^ affect *pmf* formation and dissipation (32). Higher ATP consumption in C4 species results in faster turnover of inorganic phosphate and substrate availability for ATP synthase (33), potentially resulting in altered control of ΔpH-dependent NPQ than in C3 photosynthesis.

The present work aimed to characterise differences in NPQ relaxation between C3 and C4 species using a combination of spectroscopic, chemical, and molecular approaches, including C4 CEF mutants. To account for the strong confounding effect of phylogenetic distance (34), we compared phylogenetically linked pairs of C3 and C4 species from three evolutionarily distinct genera operating three different C4 metabolic cycles.

## Results

### Differences in NPQ relaxation between C3 and C4 species

To compare NPQ between photosynthetic pathways, we selected C3 and C4 species from *Alloteropsis* (C3 *A. semialata* KWT, C4 *A. semialata* MDG), *Flaveria* (C3 *F. cronquistii*, C4 *F. bidentis*), and *Cleome* (C3 *T. hassleriana*, C4 *G. gynandra*). This experimental design minimises phylogenetic variation between each C3 and C4 pair, whilst maintaining substantial evolutionary distance between the three genera. The selected species represent both monocots (*Alloteropsis*) and dicots (*Flaveria, Cleome*), three independent C4 origins (35, 36), and the three major decarboxylating enzymes found across C4 photosynthetic species: NADP-ME/PEPCK in C4 *A. semialata MDG* (37), NADP-ME in C4 *F. bidentis* (38), and NAD-ME in C4 *G. gynandra* (39).

NPQ induction and relaxation responses were measured during a 1 hour photoperiod (600 µmol m^-2^ s^-1^ PFD) followed by 25 minutes of darkness. Since NPQ components can be resolved based on their relaxation kinetics (1, 2) and given current interest in NPQ relaxation for improving photosynthesis (40), we focused on NPQ following the light-to-dark transition (**Fig. 1A**, full traces in **Fig. S1**). NPQ relaxation was faster in all C4 species relative to their C3 counterparts, resulting in significantly lower NPQ across the dark period. Integrated NPQ across the dark recovery period was 40% lower in C4 *Alloteropsis semialata MDG* than in C3 *Alloteropsis semialata KWT*, 33% lower in C4 *Flaveria bidentis* than in C3 *Flaveria cronquistii*, and 22% lower in C4 *Gynandropsis gynandra* than in C3 *Tarenaya hassleriana* (values and statistics in **Table S1**).

**Fig. 1:**
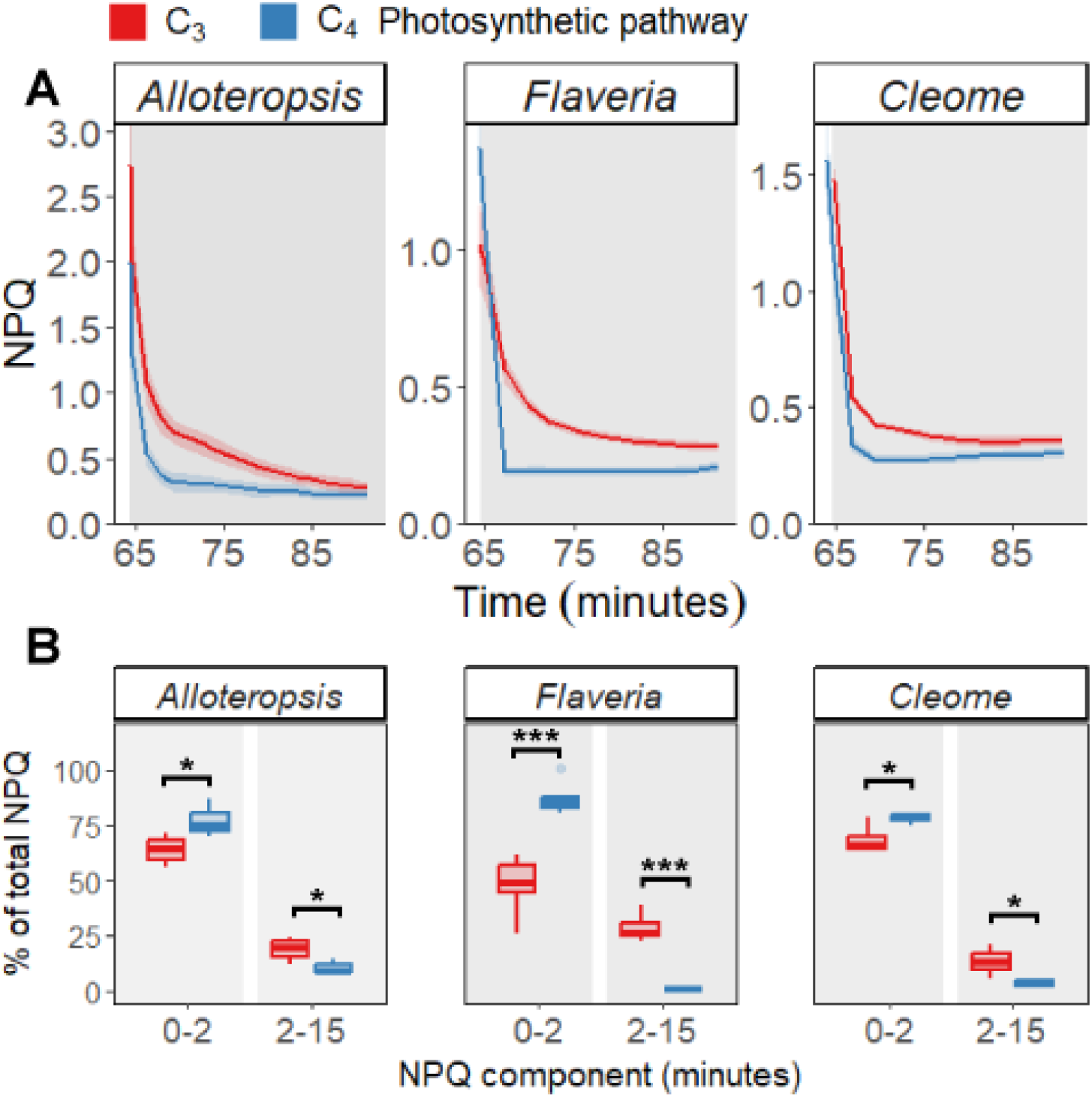
Differences in NPQ relaxation in phylogenetically linked C3 and C4 species (n=5). **A)** NPQ relaxation after 1h of illumination at 600 µmol m^-2^ s^-2^ PFD preceded by 5m of darkness for *F*_*v*_*/F*_*m*_. Ribbons represent standard error of the mean. **B)** NPQ composition based on time relaxation kinetics as a percentage of total NPQ. Asterisks indicate significant differences between C3 and C4 species found by one-way ANOVA (* *P* ≤ 0.05, ** *P* ≤ 0.01, *** *P* ≤ 0.001).

The NPQ relaxation kinetics of C3 and C4 species were underpinned by clear differences in NPQ composition (**Fig. 1B**, values and statistics in **Table S2**). NPQ components were analysed by separating NPQ relaxation into different timescales of deactivation expressed as a function of total NPQ. Fast-relaxing components (0-2 min) constituted a significantly greater proportion of NPQ in all C4 species than in their C3 pairs and conversely, slower-relaxing NPQ components (2-15 min) were a larger part of NPQ in C3 species, contributing to a more exponential decay.

### Effects of chemical inhibition of ΔpH and xanthophyll-dependent components

Fast-relaxing NPQ components include ΔpH-sensitive and xanthophyll-dependent qE in the first two minutes, and qZ dissipation in 2-15 minutes (2). To identify the elements responsible for NPQ differences between C3 and C4 species, leaves were infiltrated with nigericin to collapse the proton gradient or dithiothreitol (DTT) to inhibit the xanthophyll cycle, and integrated NPQ of treated leaves during the light period was compared to a control to assess NPQ dependence on both mechanisms (**Fig. 2**, full traces in **Fig. S2**). Nigericin infiltration resulted in significantly greater suppression of NPQ in C4 than in C3 species (**Fig. 2A**, values and statistics in **Table S3**), indicating higher reliance on ΔpH for C4 NPQ responses: 75±3% reduction in C4 *A. semialata MDG* vs. 58±9% in C3 *A. semialata KWT*, 88±2% in C4 *F. bidentis* vs. 62±4% in C3 *F. cronquistii*, and 94±3% in C4 *G. gynandra* vs. 68±3% in C3 *T. hassleriana*. Although both qE and xanthophyll de-epoxidation depend on ΔpH, the lack of significant differences in NPQ suppression by DTT in any C3 and C4 pairs (**Fig. 2B**) suggests that differences in C4 NPQ mostly stem from ΔpH-sensitive qE rather than qZ or xanthophyll-dependent qE.

**Fig. 2:**
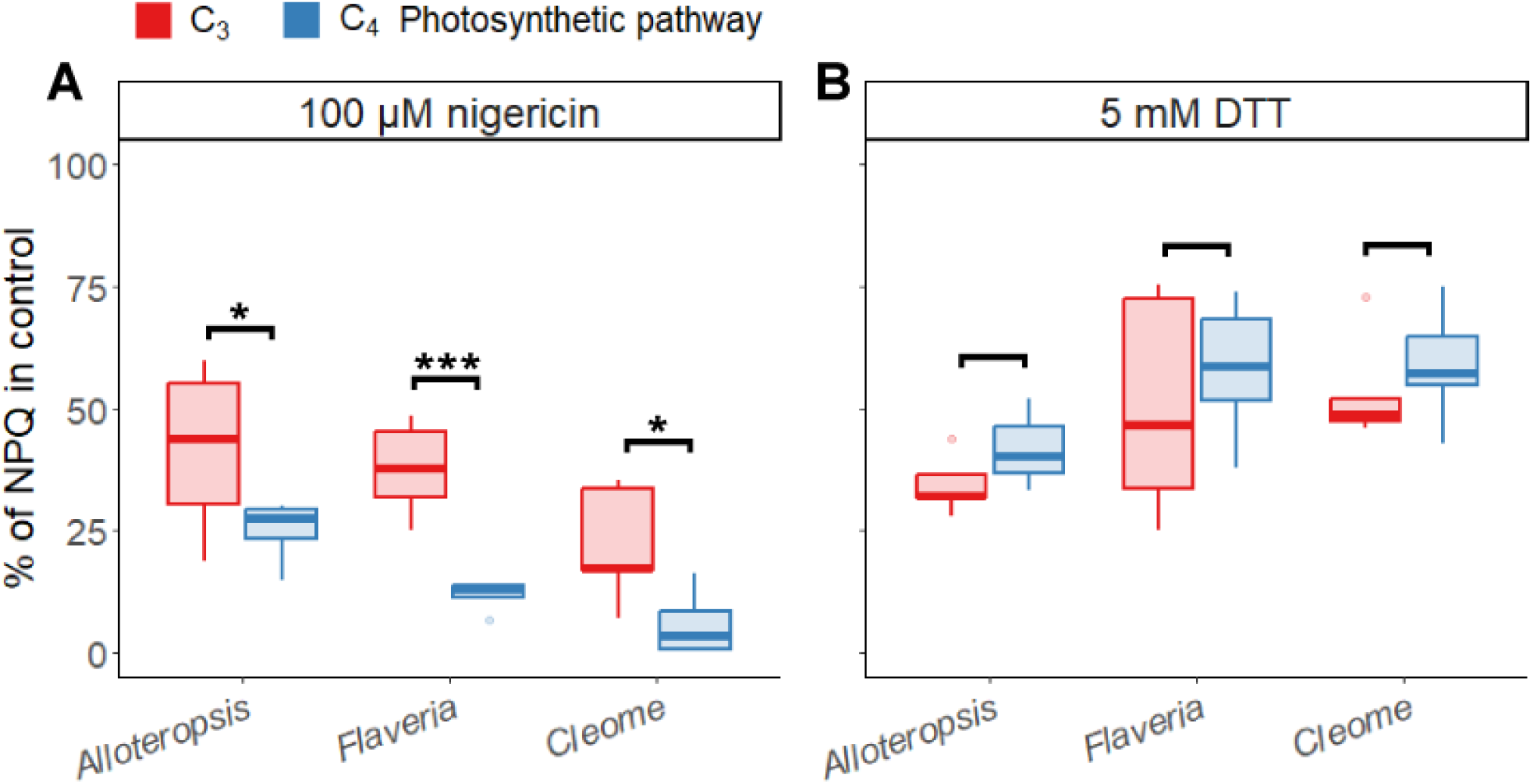
Effect of ΔpH disruption and xanthophyll cycle inhibition on C3 and C4 NPQ. AUC of NPQ induction of leaves infiltrated with **A)** 100 µM nigericin to collapse the proton gradient and **B)** 5 mM dithiothreitol to inhibit the xanthophyll cycle, as a percentage of a control (n=5). Asterisks indicate significant differences between C3 and C4 species found by one-way ANOVA (* *P* ≤ 0.05, ** *P* ≤ 0.01, *** *P* ≤ 0.001).

### Effects of the removal of photosynthetic and photorespiratory electron sinks

The higher rates of CEF to satisfy ATP:NADPH requirements (23) and larger electron sinks found in C4 photosynthesis (33) may contribute to the formation and collapse of ΔpH. To test whether fast NPQ relaxation in C4 species is linked to C4 metabolism, we repeated our NPQ measurements in 2% O_2_ + 0 ppm CO_2_ air (**Fig. 3**). The suppression of photosynthesis and photorespiration slowed down NPQ relaxation and removed differences between C3 and C4 *Alloteropsis* and *Cleome* pairs. Unlike the step-like decay observed in ambient air (**Fig. 1A**), NPQ in C4 *A. semialata MDG* and C4 *G. gynandra* followed the approximately exponential decay pattern of their C3 counterparts in 2% O_2_ + 0 ppm CO_2_ (**Fig. 3A**, values and statistics in **Table S1**), and had similar relative contributions of the 0-2 min and 2-15 min components to total NPQ (**Fig. 3B, Table S2**). In contrast, differences in NPQ kinetics between C3 and C4 species seemed enhanced in *Flaveria* in 2% O_2_ + 0 ppm CO_2_ air. Integrated NPQ was lower in C4 *F. bidentis* than in C3 *F. cronquistii*; and the 0-2 min component still represented a significantly larger proportion of NPQ in C4 *F. bidentis* than in C3 *F. cronquistii*, whilst the 2-15 min component showed the opposite trend.

**Fig. 3:**
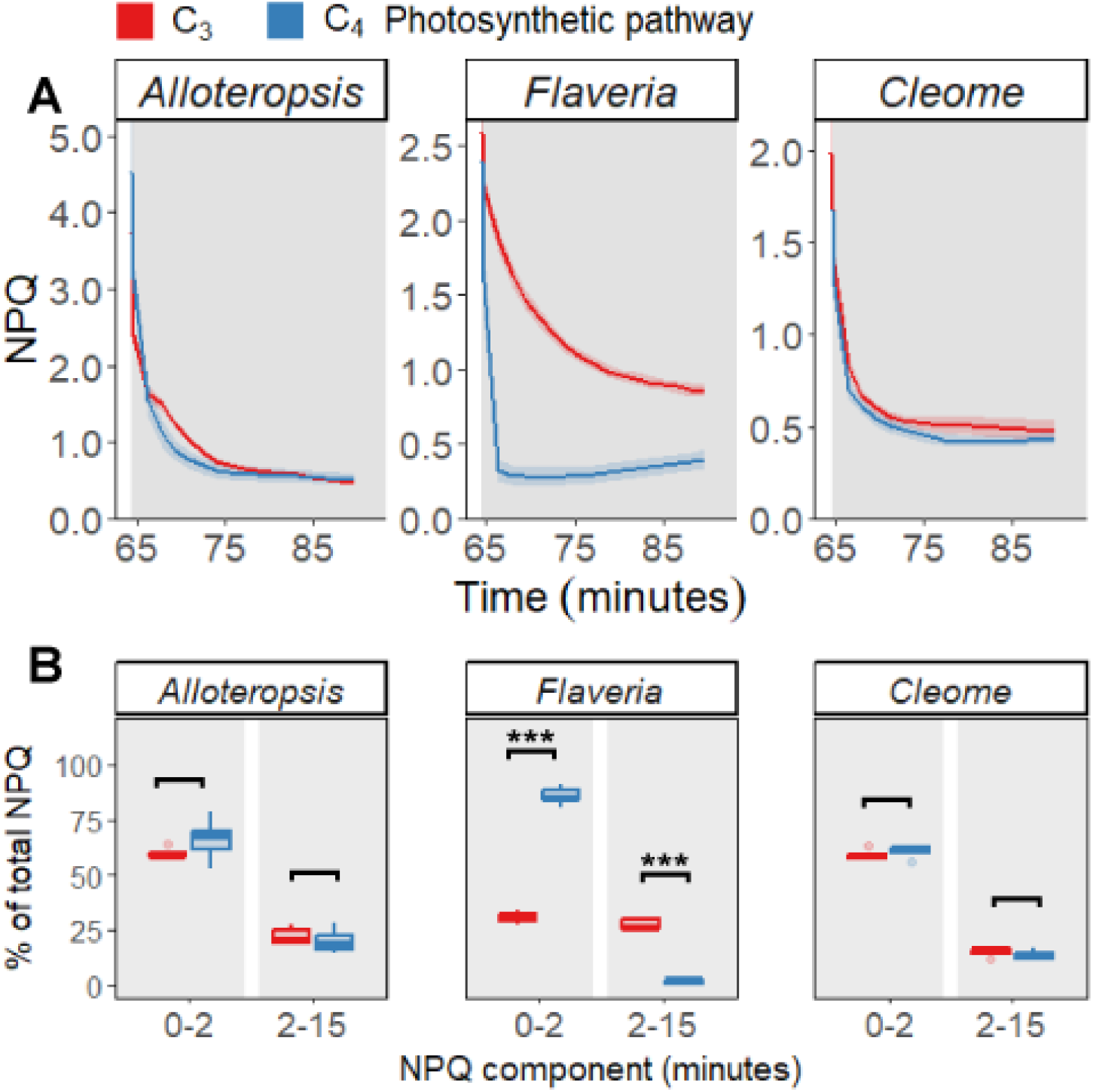
Differences in NPQ relaxation in phylogenetically linked C3 and C4 species when suppressing photosynthesis and photorespiration with 2% O_2_ and 0 ppm CO_2_ air (n=5). **A)** NPQ relaxation after 1h of illumination at 600 µmol m^-2^ s^-2^ PFD preceded by 5m of darkness for *F*_*v*_*/F*_*m*_. Ribbons represent standard error of the mean. **B)** NPQ composition based on time relaxation kinetics as a percentage of total NPQ. Asterisks indicate significant differences between C3 and C4 species found by one-way ANOVA (* *P* ≤ 0.05, ** *P* ≤ 0.01, *** *P* ≤ 0.001).

Based on the pH requirement of qE, we speculated that C4 *F. bidentis* may retain H^+^ efflux capacity in 2% O_2_ + 0 ppm CO_2_ air. We estimated gH^+^ from decay kinetics of electrochromic shift (ECS) signal measurements during brief dark intervals across 20 minutes of light to encompass the 0-2 and 2-15 min NPQ components (**Fig. 4**). Under ambient air, all C4 species tended to have higher gH^+^ than their C3 pairs (**Fig. 4A, C & E**). However, C4 *F. bidentis* retained H^+^ efflux capacity in 2% O_2_ + 0 ppm CO_2_ (**Fig. 4D**), albeit diminished compared to 21% O2 + 410 ppm CO_2_, whereas suppressing photosynthesis and photorespiration resulted in gH^+^ values approaching zero in all other species (**Fig. 4B, D & F**). These results show that fast NPQ relaxation in C4 species is strongly linked to proton efflux capacity, and that an alternative electron sink in C4 *F. bidentis* likely sustains proton efflux and qE even when photorespiration and photosynthesis are suppressed.

**Fig. 4:**
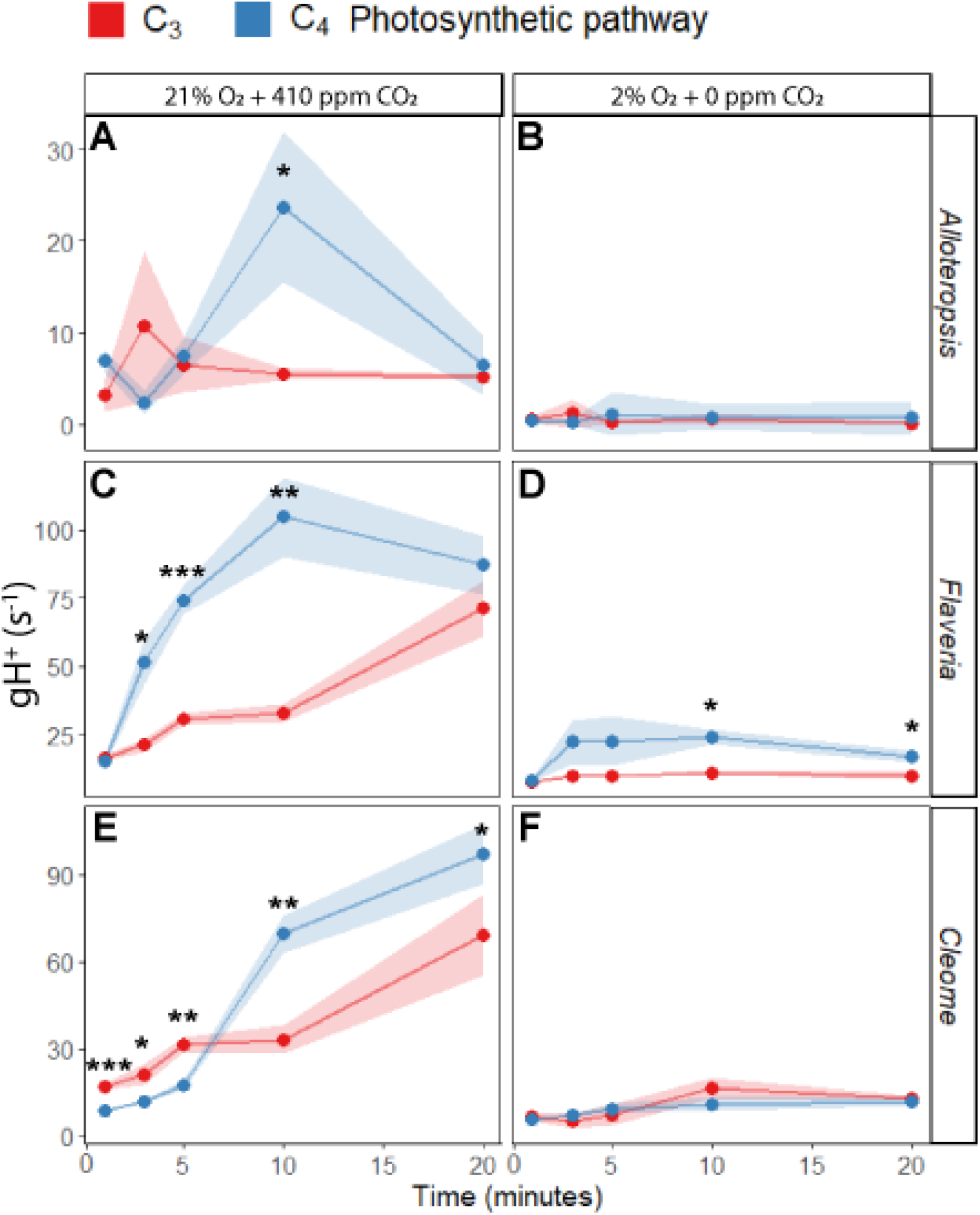
Proton conductivity of the thylakoid membrane (gH^+^) in C3 and C4 phylogenetically linked species, under ambient and 2% O_2_ air and 0 ppm CO_2_ (n=5), during 20 minutes of illumination at 600 µmol m^-2^ s^-2^ PFD. Asterisks indicate significant differences between species at a given timepoint found by one-way ANOVA (n=5, * *P* ≤ 0.05, ** *P* ≤ 0.01, *** *P* ≤ 0.001).

### Evaluating CEF in C3 and C4 species

Chlorophyll fluorescence reflects the PSII quinone redox state. In darkness, with no LEF-driven reduction, a post-illumination chlorophyll fluorescence rise (PIFR) is attributed to residual CEF transferring electrons from stromal donors to the PQ pool that then equilibrate with PSII-associated quinones (41). We measured PIFR as a proxy for CEF after 1 h light treatment, periodically flashing far-red light to preferentially excite PSI and temporarily enhance PQ oxidation (full PIFR trace in **Fig. S3**). After the far-red pulse, a larger PIFR response was observed in C3 *A. semialata KWT*, C4 *F. bidentis*, and C4 *G. gynandra* than in their phylogenetic pairs, indicating higher CEF activity in those species (**Fig. 5A**, solid lines). This is consistent with ATP:NADPH ratio requirements being higher in NADP-ME (C4 *F. bidentis*) and NAD-ME (C4 *G. gynandra*) subtypes than in mixed PEPCK pathways (C4 *A. semialata MDG*), which have lower ATP requirements (18). Leaves were also treated with PGR5/PGRL1 pathway inhibitor Antimycin A (dashed lines), which resulted in significantly lower PIFR in C3 *F. cronquistii* and C3 *T. hassleriana*, indicating a substantial contribution of PGR5/PGRL1 to PQ reduction. The addition of Antimycin A also resulted in higher PIFR values in some cases – a secondary effect of Antimycin A that has been previously attributed to additional inhibition of electron transport downstream of PQ (31). Even with this effect, these results validate existing knowledge of C3 and C4 CEF pathways– C3 species primarily use the PGR5/PGRL1 route while C4 species use both the PGR5/PGRL1 and the NDH pathways (26, 30, 31). C3 *A. semialata KWT* appears to be an exception, but given that the C3 subspecies may have reversed from a C3-C4 intermediate, CEF operation might differ from other C3 plants (42).

**Fig. 5:**
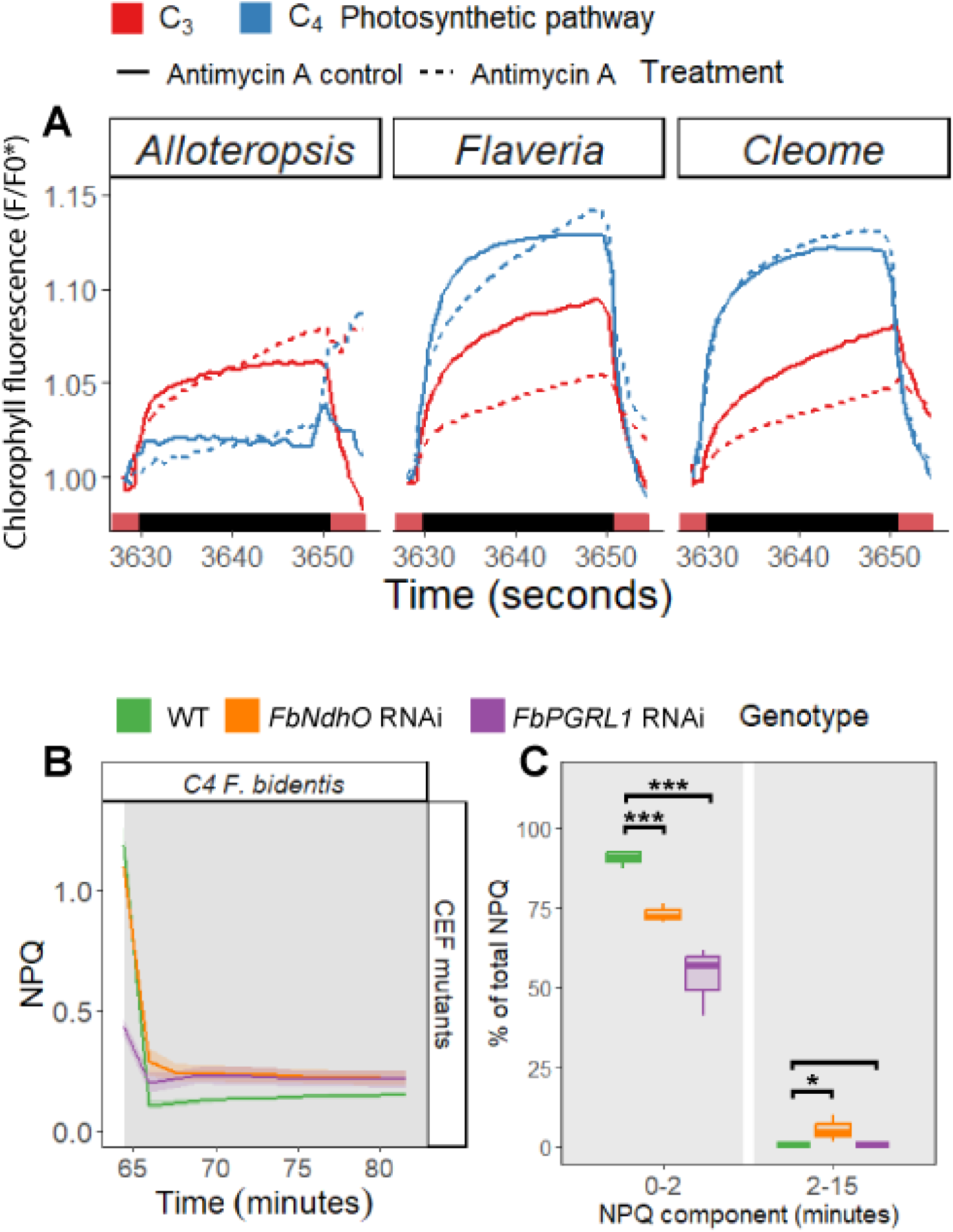
CEF in C3 and C4 phylogenetically linked species. **A)** Representative trace (n=5) of post-illumination fluorescence rise (PIFR) in leaves infiltrated with a control buffer (solid lines) or with Antimycin A (dashed lines), an inhibitor of the CEF PGR5 pathway. **B)** NPQ relaxation in C4 *F. bidentis* WT and *FbNdhO* and *FbPGRL1* RNAi knockdown mutants. Ribbons represent standard error of the mean (n=3). **C)** NPQ composition based on time relaxation kinetics as a percentage of total NPQ. Asterisks indicate significant differences between each mutant and the WT found by one-way ANOVA (n=3, * *P* ≤ 0.05, ** *P* ≤ 0.01, *** *P* ≤ 0.001).

The role of CEF in C4 NPQ relaxation was further studied in C4 *F. bidentis FbPGRL1-RNAi* and *FbNdhO-RNAi* knockdown lines (31) – as previously observed, carbon assimilation was notably lower only in *FbNdhO-RNAi* (**Fig. S4**), suggesting an impaired C4 cycle. NPQ in *FbNdhO-RNAi* decayed more slowly than the step-wise drop observed in WT and *FbPGRL1-RNAi* (**Fig. 5B**, values and statistics in **Table S4**). Whilst the 0–2 min component constituting most of WT NPQ was significantly reduced in both *FbPGRL1-RNAi* and *FbNdhO-RNAi*, the slower NPQ relaxation of *FbNdhO-RNAi* is evidenced by a more significant 2-15 min component than in WT. At the end of illumination, NPQ was lower in *FbPGRL1-RNAi* than in WT and *FbNdhO-RNAi*, suggesting that differences in NPQ composition stem from impaired relaxation in *FbNdhO-RNAi*, but from overall NPQ suppression in *FbPGRL1-RNAi*.

## Discussion

### C4 species have faster NPQ relaxation

This study sought to characterise differences in NPQ relaxation between C3 and C4 photosynthesis, by comparing phylogenetically linked *Alloteropsis, Flaveria*, and *Cleome* C3 and C4 species. Despite considerable evolutionary distance between the three tested genera, all C4 species had significantly faster and overall greater NPQ relaxation than their C3 pairs (**Fig. 1**), showing that this is likely linked to the C4 pathway. The rapid return of PSII to the unquenched state in C4 species would support higher photosynthetic quantum yields following decreases in irradiance, and could be contributing to the more sustained rates of CO_2_ assimilation that have been observed in C4 versus C3 species during light-shade transitions (43, 44). Whilst increasing the rate of photosynthetic efficiency has been remarkably successful at improving photosynthetic efficiency in C3 species (9, 10), our results suggest that this approach may result in more limited gains in carbon assimilation in C4 species given their intrinsically faster NPQ relaxation rate.

### A greater proportion of C4 NPQ is qE

The comparatively faster NPQ relaxation found in C4 plants stemmed from differences in NPQ composition between C3 and C4 species. Separation of NPQ into components based on decay timescales revealed that C4 species had a significantly higher proportion of a fast-relaxing (0-2 min) component compared to their C3 phylogenetic pairs, with C3 species exhibiting a comparatively greater proportion of NPQ relaxation within the 2-15 min timeframe (**Fig. 1B**). NPQ component qE operates within the 0-2 min timescale, responding to lumen acidification and enhanced by zeaxanthin accumulation (2, 45). When infiltrated with ΔpH-inhibitor nigericin, NPQ in C4 species was significantly more depressed than in C3 species, whereas inhibiting xanthophyll cycle activity with DTT did not result in significant differences in the extent of NPQ reduction between C3 and C4 species (**Fig. 2**). These results demonstrate that the acceleration of NPQ identified in all three C4 species involved ΔpH-dependent qE. Activation of qE by lumen pH relies on the presence of photosystem II subunit S (PsbS), which initiates the quenched LHCII state via a conformational switch upon protonation of lumen-exposed protonatable residues (46, 47). Thus, one hypothesis could be that C4 species may show slight alterations in PsbS relative abundance, structure, or interactions with LHCs to explain the enhancement of qE.

### The energetic requirements of the C4 pathway affect NPQ

When photosynthesis and photorespiration were suppressed, NPQ relaxation in C4 *A. semialata MDG* and *G. gynandra* conformed to the slower exponential decay and composition of C3 *A. semialata KWT* and *T. hassleriana* (**Fig. 3**). In ambient air, C4 metabolism largely suppresses photorespiration so these results suggest that the fast NPQ relaxation observed in *Alloteropsis* and *Cleome* C4 species is specifically related to the ATP and NADPH demand by the C4 photosynthetic pathway. qE is regulated by lumen pH, which is in turn dependent on membrane proton conductivity, gH^+^ (32). The higher gH^+^ found in all three C4 species in ambient conditions (**Fig. 4A, C & E**) could contribute to accelerated relaxation of pH-dependent NPQ, primarily qE.

Different C4 pathways have distinct ATP:NADPH requirements: in NADP-ME/PEPCK subtypes (C4 *A. semialata MDG*), the energetic balance is comparable to C3 species, whereas NADP-ME (*F. bidentis*) and NAD-ME (*G. gynandra*) pathways have elevated ATP:NADPH demands. The increased ATP demand is localised in BS cells in NADP-ME subtypes but in M cells in NAD-ME subtypes (16-18). M cells are positioned in the outer layer of Kranz anatomy, and therefore represent the majority of the chlorophyll fluorescence signals used to determine NPQ. However, in all three C4 species, the large metabolic pools of C4 cycle intermediates required to sustain diffusion gradients between M and BS cells may underpin the observed increases in gH^+^ relative to C3 species. The high gH^+^ indicates upregulated proton efflux, be it ATP synthase dependent or independent as photoprotective ΔpH has been found to be linked both to ATP synthase regulation (32, 48) as well as to thylakoid antiporters like KEA3 (49), both mechanisms that could differ in C4 photosynthesis compared to C3. Uniquely, in C4 *F. bidentis* NPQ relaxation was not slowed by the suppression of photosynthesis and photorespiration (**Fig. 3**), and gH^+^ was much higher than in the other C4 species (**Fig. 4D**), indicating the presence of an alternative electron sink. The identity of this electron sink remains unclear but it seems plausible that it relates to specific attributes of the canonical NADP-ME C4 pathway, such as the significantly larger malate pool sizes that could go towards ATP-consuming metabolic reactions (50), although sustained leakiness of the membrane via ATP synthase-dependent and independent mechanisms is also possible.

### Contribution of CEF pathways to faster NPQ in C4 species

In C3 species, CEF via the PGR5/PGRL1 pathway is essential for photoprotection by contributing to ΔpH formation and qE (24, 25). Relative to the ancestral C3 pathway, PGR5/PGRL1 abundance increases in C4 species in both M and BS cells (17, 51), whilst NDH differentially accumulates per ATP requirements: in M cells for NAD-ME pathways and in BS for NADP-ME (16, 18). Our results point to both NDH and PGR5/PGRL1 pathways contributing to fast relaxation in C4 NPQ. Both C4 *FbNdhO-RNAi* and *FbPGRL1-RNAi* lines showed altered NPQ relaxation kinetics relative to WT (**Fig. 5B**). Consistent with our findings, previous work found the PGR5/PGRL1 pathway to significantly contribute to NPQ induction in C4 *F. bidentis* (31). Here, we also show that knocking down PGRL1 results in a specific decrease in qE activation (**Fig. 5C**), probably due to deficient *pmf* formation. In contrast, reduction of NDH expression primarily slowed down NPQ responses. C4 *F. bidentis* NDH is primarily found in the BS, accounting for most of BS CEF (30). Although the chlorophyll fluorescence signal primarily comes from the M, an impaired 0-2 min NPQ component was still observed in *FbNdhO-RNAi* mutants (**Fig 5C**). The tissue-specific expression of NDH in C4 species and the deleterious effect of its suppression on growth and carbon assimilation suggests NDH is the major route for ATP provision to C4 metabolism (30, 31). Accordingly, the rate of carbon assimilation in *FbNdhO-RNAi* was greatly reduced relative to wild type (**Fig S4**). An impaired C4 cycle in *FbNdhO-RNAi* would alter inorganic phosphate availability and LEF:CEF energy balance, potentially diminishing CEF-related qE and leading to parallel decreases in gH^+^ and ΔpH in M cells (32, 33). Since the role of NDH in C4 photosynthesis has thus far only been studied in NADP-ME plants with NDH-enriched BS, it remains unclear if NDH-mediated CEF generally also plays a photoprotective role similar to PGR5/PGRL1. Future research could test this in NAD-ME species with NDH-enriched M (22), such as C4 *G. gynandra*.

The complexity of NPQ molecular mechanisms, evolution of CEF pathways, and biochemical diversity within C4 species leave many outstanding questions with relation to C4 NPQ. Nevertheless, the results presented here demonstrate that the higher rates of CEF and larger electron sinks of C4 metabolism result in enhancement of the qE component, leading to faster relaxation kinetics. The increased contribution of qE to total NPQ in C4 species mimics the engineering attempts to accelerate NPQ responses in C3 crops via single overexpression of PsbS (52, 53) or in combination with violaxanthin de-epoxidase and zeaxanthin epoxidase (9, 10), engineering attempts which may have less scope to further accelerate NPQ responses and photosynthetic efficiency in C4 species.

## Materials and Methods

### Plant material and growth conditions

C4 *F. bidentis*, C3 *F. cronquistii*, C4 *G. gynandra* and C3 *T. hassleriana* were grown in soil under growth chamber conditions at 20ºC and 150 µmol m^-2^ s^-1^ PFD over a 16-hour photoperiod. *A. semialata MDG* and *A. semialata KWT* were grown in a 4:1 mix of soil and vermiculite under semi-controlled glasshouse conditions at 18-25 ºC with supplemental lightning provided to a minimum of 140-160 µmol m^-2^ s^-1^ PFD over a 16-hour photoperiod (further detail in (54)). Measurements were conducted on fully expanded leaves during vegetative stage: at 8-10 weeks for both *Flaveria* species and *G. gynandra*, 4-6 weeks for *T. hassleriana*, and 2 weeks after vegetative propagation for both *Alloteropsis* species. C4 *F. bidentis* WT, *FbPGRL1-RNAi* and *FbNdhO-RNAi* RNAi plants with expression ∼10% of WT (31), were grown in a 3:2 mix of soil and vermiculate, in a growth chamber at 24ºC and 250 µmol m^-2^ s^-1^ PFD over a 12-hour photoperiod. Young, fully-expanded *FbNdhO-RNAi* leaves were measured after 12-16 weeks and of all other plants after 8-10 weeks.

### Chlorophyll fluorescence measurements

Chlorophyll fluorescence was measured with a gas exchange system (LI-6400XT, LI-COR, Lincoln, NE, USA) equipped with a leaf chamber fluorometer (6400-40 LCF, LI-COR). Chamber conditions were controlled at 410 or 0 ppm sample CO_2_ concentration (the latter with 2% O_2_ air), 40-60% relative humidity, 25ºC block temperature, 300 µmol s^-1^ flow rate, and 10% blue (470 nm) and 90% red actinic light (630 nm). The LCF used a 0.25 Hz modulated measuring light and a multiphase flash (55) to measure chlorophyll fluorescence parameters.

Leaves were dark-adapted until stomatal conductance and net CO_2_ exchange rate stabilised (30-60 min), illuminated with 600 µmol m^-2^ s^-1^ PFD for 1 hour, and returned to darkness for 25 minutes. Multiphase saturating flashes at 4000 µmol m^-2^ s^-1^ PFD were used to measure steady and maximal fluorescence after dark-adaption (*F* and *F*_*m*_), and during photoperiod and dark recovery (*F’* and *F*_*m*_*’*), occurring five minutes before illumination (for *F*_*v*_*/F*_*m*_); after 3, 5, 10, 15, 25, 35, 45, and 60 minutes of light exposure; and 30s after dark transition, then every 90s thereafter. NPQ was derived from fluorescence measurements (56). NPQ relaxation was compared by calculating integrated NPQ from area under the curve (AUC); and 0–2 min and 2–15 min component contributions were expressed as the integrated NPQ within each phase relative to total post-illumination NPQ.

Chlorophyll fluorescence rise during dark recovery (following (41)) was monitored for 2.5 minutes in darkness after the photoperiod, interspersed by 5s of far-red (FR) light to oxidise the PQ pool (740 nm, ∼50 μmol of photons m^−2^ s^−1^) every 20-25s from instrument variation. We compared the PIFR of the dark period after FR, normalised to the final fluorescence value post-oxidation (*F0\**, see **Fig. S3**). Representative data from five biological replicates of the subsequent rise in fluorescence is presented.

### Nigericin, DTT, and Antimycin A leaf infiltration

Dark adapted leaves were vacuum infiltrated with NPQ inhibitors in a syringe with a buffer (20 mM HEPES/KOH pH 7.0) supplemented with either 100 µM nigericin, 5 mM DTT, or 250 µM Antimycin A. Controls were infiltrated with buffer and equivalent volume of solvent without inhibitor. The effect of NPQ inhibitors was estimated by calculating NPQ AUC during the light period and expressing as a proportion of control NPQ AUC (**Fig S2** for full NPQ traces).

### Electrochromic Shift measurements

The ECS signal was measured as absorptance changes at 515 nm using 535 nm as an isosbestic waveband (Dual-KLAS fitted with a P515/535 module, Heinz Walz GmbH, Effeltrich, DE) (57). The fore-optics were integrated in a custom measuring chamber (GFS-3000 measuring chamber for DUAL-KLAS, Walz) with temperature controlled at 25^°^C. Chamber conditions were controlled via the console of a LI-6800 gas exchange system (LI-COR, USA) at 410 or 0 ppm sample CO_2_ concentration (the latter with 2% O_2_ air), 60% relative humidity, and 200 µmol s^-1^ flow rate. Leaves were dark-adapted, and illuminated with 600 µmol m^-2^ s^-1^ PFD for 20 minutes. Dark Interval Relaxation Kinetics (DIRK) measurements of the ECS signal (58) were taken at 1, 3, 5, 10, and 20 minutes of illumination. Estimates of thylakoid membrane proton conductivity (gH^+^) were obtained from the inverse of the decay time constant (τ_ECS_) of a single exponential decay fitted to the first 300 milliseconds of the dark interval (59).

### Gas concentration manipulation

To suppress photosynthesis and photorespiration, a pre-mixed 2% O_2_ and 98% N_2_ gas mixture (BOC Ltd., Woking, UK) was supplied to the LI-6400XT or LI-6800 using a mass flow controller (EL-FLOW, Bronkhorst Hight-tech BV, Ruurlo, NL), and CO_2_ controlled at 0 ppm.

### Statistical analysis

One-way ANOVA was conducted on independent experiments comparing each C3 and C4 phylogenetic pair, and comparing *FbPGRL1-RNAi* and *FbNdhO-RNAi* mutants with WT. Assumptions of normality and homogeneity of variance were tested for, respectively with a Shapiro-Wilk and Bartlett’s test.

## Acknowledgments and funding sources

For providing the original plant material, we thank Dr. Luke Dunning (*Alloteropsis* cuttings), Dr. Marjorie Lundgren (*F. cronquistii* cuttings), Prof. Peter Westhoff (*F. bidentis* seeds) and Prof. Julian Hibberd (*T. hassleriana* seeds). Additional thanks to Prof. Hibberd and Prof. David M. Kramer for kindly consulting on our results, and to Dr. Gustaf Degen for his helpful guidance on ECS.

LAC was jointly funded by the Cambridge Trust; and by Consejo Nacional de Ciencia y Tecnología (CONACyT). This work was supported by the Biotechnology and Biological Sciences Research Council (BBSRC) via grant BB/T007583/1 awarded to JK. For the purpose of open access, the authors have applied a Creative Commons Attribution (CC BY) licence to any Author Accepted Manuscript version arising from this submission.

## Author Contributions

JK and LAC conceived the study and designed the experiments. AN measured the CEF *F. bidentis* mutants developed by YM. LAC carried out all other experiments, data analysis and interpretation, and wrote the manuscript. RLV helped with the 2% O_2_ experimental setup and provided support with gas exchange experiments. JW and CRGS helped with developing ECS protocols, and JW with chemical infiltration protocols. ELB procured the initial plant material. All authors contributed to and reviewed the final manuscript. Descriptive statistics and figures were created using R 4.1.1 on RStudio 2023.03.1+446.

## Supporting Information

Fig. S1: NPQ induction and relaxation in phylogenetic pairs

Fig. S2: NPQ induction and relaxation with chemical inhibitors

Fig. S3: Sample full trace of Post-Illumination Fluorescence Rise

Fig. S4: Carbon assimilation of CEF mutants

Table S1: Values and statistics of integrated NPQ across dark recovery in phylogenetic pairs

Table S2: Values and statistics of NPQ composition in phylogenetic pairs

Table S3: Values and statistics of NPQ with chemical inhibitors in phylogenetic pairs

Table S4: Values and statistics of NPQ composition in CEF mutants

