## Supplemental Tables S1-4 and Figs S1-4 for "Faster relaxation of nonphotochemical quenching (NPQ) in C4 than in C3 species"

**Table S1: Values and statistics of integrated NPQ across dark recovery in phylogenetic pairs.** Integrated NPQ and standard error of the mean across dark recovery period in C3 and C4 species during the 25 minutes of darkness following 1h of illumination at 600 µmol m^-2^ s^-2^ PFD, in ambient and 2% O_2_ and 0 ppm CO_2_ air (n=5). One-way ANOVA was conducted between phylogenetically linked species (degrees of freedom; *F-*value; *p-*value), significant *p*-values are shown in bold.

| **Experiment** | **Species** | **Integrated NPQ** | **ANOVA** |
| --- | --- | --- | --- |
| Ambient conditions | C4 *A. semialata MDG* | 11.59±1.50 | 1,8; 5.85; **0.049** |
|  | C3 *A. semialata KWT* | 19.21±2.54 |  |
|  | C4 *F. bidentis* | 8.35±0.68 | 1,8; 17.72; **0.003** |
|  | C3 *F. cronquistii* | 12.45±0.73 |  |
|  | C4 *G. gynandra* | 11.63±.96 | 1,8; 6.14; **0.042** |
|  | C3 *T. hassleriana* | 14.86±0.74 |  |
| 2% O_2_ + 0 ppm CO_2_ | C4 *A. semialata MDG* | 22.29±3.06 | 1,8; 0.225; 0.65 |
|  | C3 *A. semialata KWT* | 23.81±0.99 |  |
|  | C4 *F. bidentis* | 10.93±1.90 | 1,8; 87.09; **≤0.001** |
|  | C3 *F. cronquistii* | 32.81±1.38 |  |
|  | C4 *G. gynandra* | 14.39±1.01 | 1,8; 1.26; 0.28 |
|  | C3 *T. hassleriana* | 16.31±1.39 |  |

**Table S2: Values and statistics of NPQ composition in phylogenetic pairs.** NPQ composition based on time relaxation kinetics as a percentage of total NPQ with standard error of the mean in C3 and C4 species, in ambient and 2% O_2_ and 0 ppm CO_2_ air (n=5). One-way ANOVA was conducted between phylogenetically linked species for each component (degrees of freedom; *F-*value; *p-*value), significant *p*-values are shown in bold.

| **Experiment** | **Component** | **Species** | **Component of total NPQ (%)** | **ANOVA** |
| --- | --- | --- | --- | --- |
| Ambient conditions | 0-2 minutes | C4 *A. semialata MDG* | 77±3 | 1,8; 5.77; **0.047** |
|  |  | C3 *A. semialata KWT* | 64±3 |  |
|  |  | C4 *F. bidentis* | 87±3 | 1,8; 31.58; **≤ 0.001** |
|  |  | C3 *F. cronquistii* | 47±6 |  |
|  |  | C4 *G. gynandra* | 78±1 | 1,8; 7.56; **0.028** |
|  |  | C3 *T. hassleriana* | 68±3 |  |
|  | 2-15 minutes | C4 *A. semialata MDG* | 10±2 | 1,8; 5.80; **0.049** |
|  |  | C3 *A. semialata KWT* | 19±2 |  |
|  |  | C4 *F. bidentis* | 0±0 | 1,8; 98.65; **≤0.001** |
|  |  | C3 *F. cronquistii* | 28±3 |  |
|  |  | C4 *G. gynandra* | 3±1 | 1;8, 11.31; **0.012** |
|  |  | C3 *T. hassleriana* | 13±3 |  |
| 2% O_2_ + 0 ppm CO_2_ | 0-2 minutes | C4 *A. semialata MDG* | 66±4 | 1,8; 4.01; 0.061 |
|  |  | C3 *A. semialata KWT* | 60±1 |  |
|  |  | C4 *F. bidentis* | 86±2 | 1,8; 4.72; **≤0.001** |
|  |  | C3 *F. cronquistii* | 31±1 |  |
|  |  | C4 *G. gynandra* | 60±1 | 1,8; 2.39; 0.14 |
|  |  | C3 *T. hassleriana* | 59±1 |  |
|  | 2-15 minutes | C4 *A. semialata MDG* | 20±2 | 1,8; 1.92; 0.18 |
|  |  | C3 *A. semialata KWT* | 22±2 |  |
|  |  | C4 *F. bidentis* | 2±1 | 1,8; 357.03; **≤0.001** |
|  |  | C3 *F. cronquistii* | 27±1 |  |
|  |  | C4 *G. gynandra* | 13±1 | 1,8; 2.51; 0.13 |
|  |  | C3 *T. hassleriana* | 15±1 |  |

**Table S3: Values and statistics of NPQ with chemical inhibitors in phylogenetic pairs.** Integrated NPQ and standard error of the mean in C3 and C4 species infiltrated with 100 µM nigericin to collapse the proton gradient or 5 mM dithiothreitol to inhibit the xanthophyll cycle, as a percentage of a control across 1h of illumination at 600 µmol m^-2^ s^-2^ PFD (n=5). One-way ANOVA was conducted between phylogenetically linked species (degrees of freedom; *F-*value; *p-*value), significant *p*-values are shown in bold.

| **Inhibitor** | **Species** | **Integrated NPQ relative to control (%)** | **ANOVA** |
| --- | --- | --- | --- |
| 100 µmol Nigericin | C4 *A. semialata MDG* | 25±3 | 1,8; 5.52; **0.05** |
|  | C3 *A. semialata KWT* | 42±9 |  |
|  | C4 *F. bidentis* | 12±2 | 1,8; 26.31; **≤0.001** |
|  | C3 *F. cronquistii* | 38±4 |  |
|  | C4 *G. gynandra* | 6±3 | 1,8; 6.80; **0.03** |
|  | C3 *T. hassleriana* | 22±6 |  |
| 5 mM DTT | C4 *A. semialata MDG* | 42±5 | 1,8; 1.90; 0.22 |
|  | C3 *A. semialata KWT* | 34±3 |  |
|  | C4 *F. bidentis* | 59±6 | 1,8; 0.39; 0.55 |
|  | C3 *F. cronquistii* | 51±10 |  |
|  | C4 *G. gynandra* | 59±5 | 1,8; 0.56; 0.47 |
|  | C3 *T. hassleriana* | 53±5 |  |

**Table S4: Values and statistics of NPQ composition in CEF mutants.** NPQ composition based on time relaxation kinetics as a percentage of total NPQ with standard error of the mean in C4 *F. bidentis* CEF mutants (n=3). One-way ANOVA was conducted between each mutant and the wild type for each component (degrees of freedom; *F-*value; *p-*value), significant *p*-values are shown in bold.

| **Component** | **Genotype** | **Component of total NPQ (%)** | **ANOVA with WT** |
| --- | --- | --- | --- |
| 0-2 minutes | WT | 90±1 | // |
|  | *FbNndhO* RNAi | 73±2 | 1,6; 94.26; **≤0.001** |
|  | *FbPGRL1* RNAi | 53±6 | 1,6; 61.46; **≤0.001** |
| 2-15 minutes | WT | -3±1.2 | // |
|  | *FbNndhO* RNAi | 5±2 | 1,6; 13.85; **0.01** |
|  | *FbPGRL1* RNAi | -3±2 | 1,6; 0.09; 0.78 |

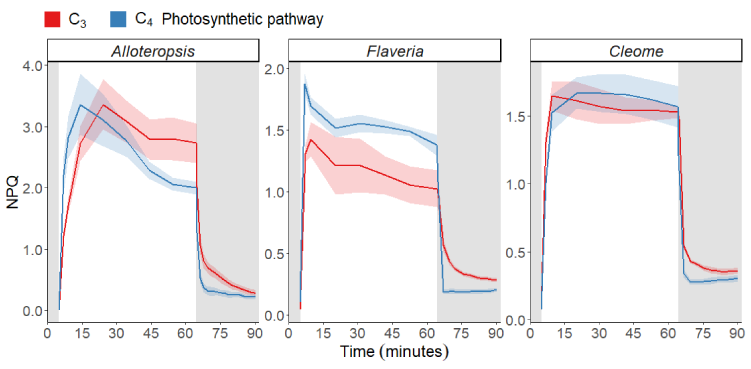
**Fig. S1: NPQ induction and relaxation in phylogenetic pairs.** NPQ induction and relaxation in phylogenetically linked C3 and C4 species during 1h of illumination at 600 µmol m^-2^ s^-2^ PFD preceded by 5m and followed by 25m of darkness. Ribbons represent standard error of the mean (n=5).

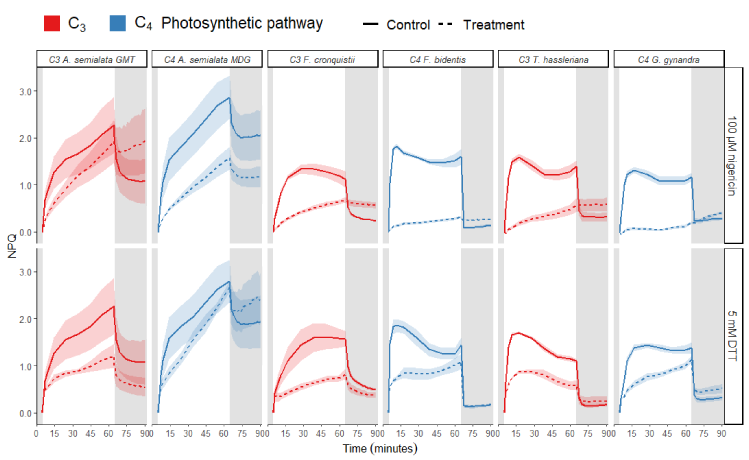
**Fig. S2: NPQ induction and relaxation with chemical inhibitors.** NPQ induction and relaxation during 1h of illumination at 600 µmol m^-2^ s^-2^ PFD preceded by 5m and followed by 25m of darkness in C3 and C4 species infiltrated with a control buffer (solid lines); 100 µM nigericin to collapse the proton gradient, or 5 mM dithiothreitol to inhibit the xanthophyll cycle (dashed lines). Ribbons represent standard error of the mean (n=5).

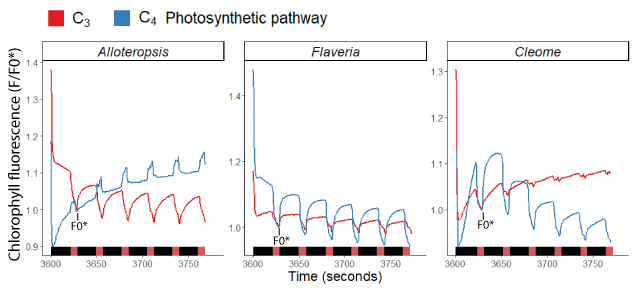
**Fig. S3: Sample full trace of Post-Illumination Fluorescence Rise.** Representative full traces of Post Illumination Fluorescence Rise measurements in C3 and C4 phylogenetically paired species after 1h of illumination at 600 µmol m^-2^ s^-2^ PFD. F’ was measured during 20-25s dark intervals from instrument variation interspersed by 5s of far-red light. The fluorescence trace was normalised to *F0**, the final fluorescence value after the first pulse of far-red light (indicated in figure).

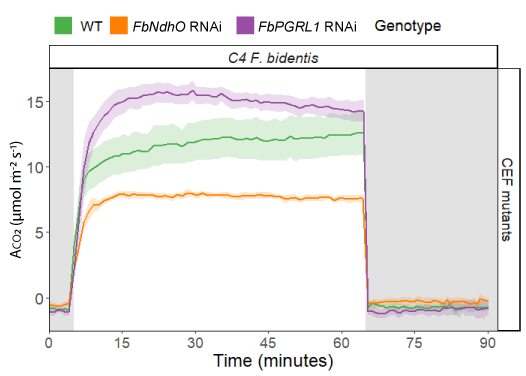
**Fig. S4: Carbon assimilation of CEF mutants.** Carbon assimilation in WT and CEF mutants of C4 *F. bidentis* during 1h of illumination at 600 µmol m^-2^ s^-2^ PFD preceded by 5m and followed by 25m of darkness. Ribbons represent standard error of the mean (n=3).
